# Predicting the structure and function of coalesced microbial communities

**DOI:** 10.1101/101436

**Authors:** Pawel Sierocinski, Kim Milferstedt, Florian Bayer, Tobias Großkopf, Mark Alston, Sarah Bastkowski, David Swarbreck, Phil J Hobbs, Orkun S Soyer, Jérôme Hamelin, Angus Buckling

## Abstract

Microbial communities commonly coalesce in nature, but the consequences for resultant community structure and function is unclear. Consistent with recent theory, we demonstrate using methanogenic communities that the most productive communities in isolation dominated when communities were mixed. As a corollary of this dynamic, total methane production increased with the number of inoculated communities. The cohesion and dominance of single communities was explained by more “niche-packed” communities being both more efficient at exploiting resources and resistant to invasion, rather than a function of the average performance of component species. These results are likely to be relevant to the ecological dynamics of natural microbial communities, as well as demonstrating a simple method to predictably enhance microbial community function in biotechnology, health and agriculture.

---

Immigration has major impacts on both the structure and function of microbial communities(*1*, *2*) and evolutionary dynamics of populations(*3*). While most work on immigration in microbial ecology deals with relatively low numbers and diversity of immigrants, this does not capture the natural context, which frequently involves the coalescence of entire communities(*4*, *5*). The consequences, if any, of such community coalescence are unclear, although existing theoretical(*6*–*9*) and empirical(*10*–*13*) studies suggest coalescence can lead to a single community dominating the mixture, rather than a more chimeric outcome. A recent extension(*9*) of ecological theory(*14*–*17*) suggests that this dominance can be predicted from how completely communities exploit diverse resources in the environment, as these “niche-packed” communities will be more cohesive and harder to invade. We test this prediction, and its corollary that coalescing communities should increase productivity, using complex anaerobic microbial communities, for which methane production is a measure of community resource use efficiency (*18*).

Anaerobic digestion is a multi-stage process carried out by highly diverse bacterial and archaeal communities. Methanogenesis is the final stage of the process and results from the conversion of H_2_, CO_2_ and short chain fatty acids produced by hydrolysis and fermentation of more complex organic material(*18*). It is carried out exclusively by methanogenic Archaea and is the only thermodynamically feasible way of actively removing inhibitory end-metabolites under the conditions where anaerobic digestion occurs(*18*). Methane production can therefore be a useful proxy of the ability of an anaerobic community to fully exploit available resources, and should correlate with community-level productivity. Moreover, methanogenic communities are characterized by complex cross-feeding interactions (*19*), and hence the importance of niche-partitioning in shaping community performance is likely to be particularly important (9). As such, methanogenic communities provide a useful and relevant system to investigate the interplay between community productivity and community coalescence.

We first determined the methane production and composition of two natural methanogenic communities grown in isolation or as a mixture in laboratory scale Anaerobic Digestors (ADs) over 5 weeks. To remove any confounding effects caused by differences in starting density of tested communities, we standardized microbial density based on qPCR-estimated counts of 16S rDNA copies. We found that the methane production of the mixed community was initially intermediate between the two individual communities, but soon started to produce gas at a rate indistinguishable from the more productive of the individual communities (Figure 1A). The composition of the mixed community at the end of the experiment most closely resembled the best performing individual community (Figure 1B). These results suggest that the most productive community dominates a mixed community, thus enhancing productivity of the mixed community beyond the average of its individual community components.

**Figure 1:**
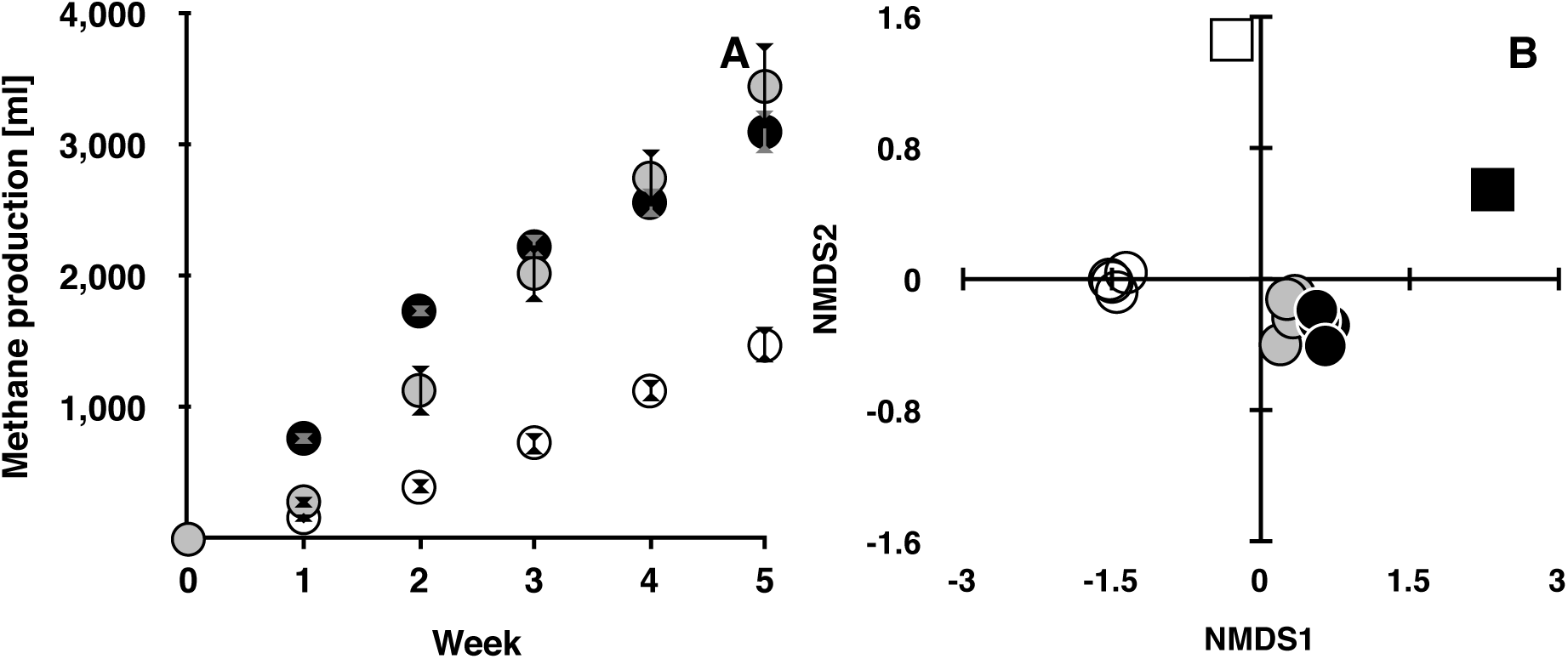
Methane production predicts the composition when two communities coalesce. A) Cumulative methane production in ml (±SEM) over time of: community P01 (white circles), community P05 (black circles) and their mixes (grey circles). Cumulative methane production differed between treatments (*F*_2,9_ = 23.2, *P* < 0.001), but did not differ between the mixed community and P05 (Tukey-Kramer HSD: *P* = 0.5). P01 was lower than both other treatments (*P* < 0.001 in both cases). B) NMDS plot of unweighted UniFrac of communities P01 (white), P05 (black) and their mixes (grey). Ancestral samples are represented by squares with samples from the endpoint of the experiment by circles. At the endpoint, P05 was compositionally more similar to the mixtures than P01, based on both unweighted (mean distance to each mixture for each replicate single community: *t*_6_ = 8.3, *P* < 0.001) and weighted (*t*_6_ = 2.3, *P* = 0.03) UniFrac distances.

We next investigated if individual communities’ methane production can predict dominance when multiple communities are mixed. Consistent with the results from two communities, methane production in mixtures of ten communities was higher than the average of the individual communities, but did not differ from the best performing single community (Figure 2A). Moreover, the community composition of mixtures (which varied little between replicates) most closely resembled the highest performing individual community (Figure 2B). More generally, the more similar was an individual community’s composition to the mixtures, the higher was its methane production (Figure 2C).

**Figure 2:**
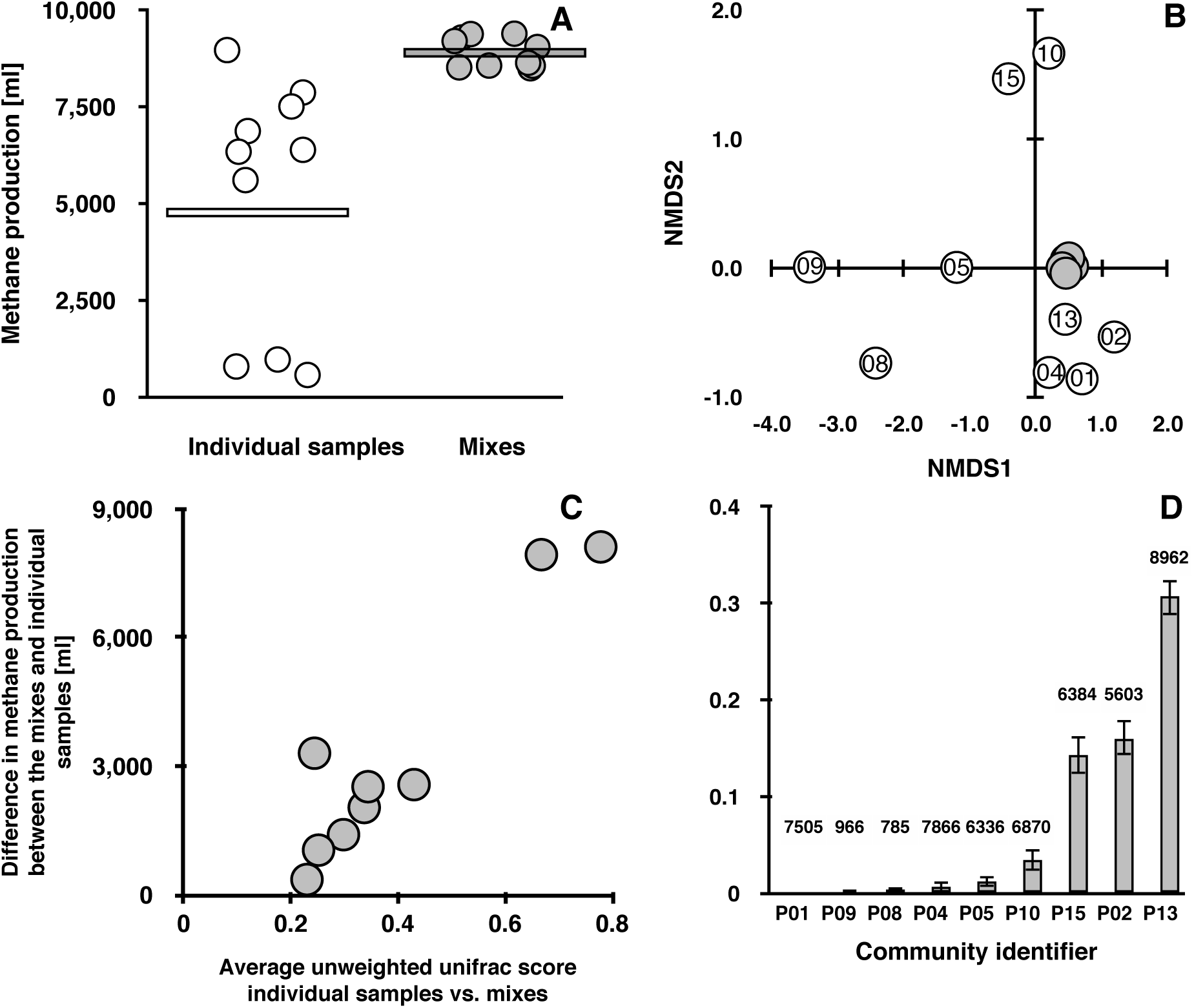
Methane production predicts the greatest contributor to coalesced community composition. A) Cumulated methane production of Mixed (grey) and Individual communities (white). Average performance shown as a horizontal line. Mean cumulative methane production was greater for mixtures than for individual communities (*t*-test: *P* < 0.001 in 9 cases), except community P13 (best performer). B) NMDS plot of unweighted unifrac of 10 mixtures (grey) and 9 individual communities (white). Numbers in circles refer to individual community identifiers (Table 1). Community P13 was significantly closer in composition to the 10 mixed communities than any other community (weighted and unweighted UniFrac distances; Paired *t*-tests; *P* < 0.001, in all cases). Note: DNA yield from community P06 was insufficient for sequencing, therefore it is excluded from this and following graphs. C) Difference between the average methane production of the mixes and individual sample and community composition according to unweighted UniFrac (Spearman ρ = 0.86, P < 0.001)). Same correlation stands for unweighted UniFrac (Spearman ρ = 0.75, P < 0.02). D) Estimated contribution of each individual community towards the 10 coalesced communities (±SEM). Values over the bars indicate methane production of each individual community over the course of the experiment [ml].

The hypothesized mechanism underlying these results is that interactions within a community determine both its productivity and cohesion. Note that this hypothesis does not invoke any “higher order selection” resulting in community-level adaptation(*16*), but that communities whose members have evolved to exploit ecological niches more completely or more efficiently (more “niche-packed” communities) are both more productive and can be less readily invaded(*9*,*17*,*20*). An alternative explanation is that community productivity is simply a function of the performance of individual species, and the most productive community dominates because it members are on average are better at exploiting resources than their ecological counterparts in other communities. If the latter is true, then community productivity will not only predict which community dominates, but also the relative contribution each individual community makes to the mixture. Specifically, species in the second most productive community will have the second highest performance on average, and hence should contribute to the mix of communities more than the third most productive community, and so on.

To distinguish between these hypotheses, we used a non-negative least squares (NNLS) approach to estimate the actual contribution of each community to the mixtures, rather than just the similarity between the single and mixed communities, as above. This confirmed that the most productive community dominated the mixture (Figure 2D), but crucially there was not a monotonic relationship between methane production and contribution to the mixture across the other communities. Indeed, the second and third most productive communities (notably communities P01 and P04) were greatly under-represented in the final mixture. This under-representation of such productive communities is entirely consistent with niche-packing determining community success: these communities were very similar in composition to the most productive community and hence presumably could not occupy the same ecological niches in the mixed community (Figure 2D). Further lines of evidence support the key role of community level properties in explaining our results. First, within-community diversity positively correlated with methane production – a finding consistent with greater niche packing(*14*) (Figure 3A). Second, the density of the organisms directly responsible for methane production, the methanogenic Archaea, did not correlate with methane production (Figure 3B), emphasizing the importance of interactions with other taxa in determining methane production. This is in contrast to the density of bacteria (Figure 3C), the majority of organisms in methanogenic communities, which positively correlated with methane production.

**Figure 3:**
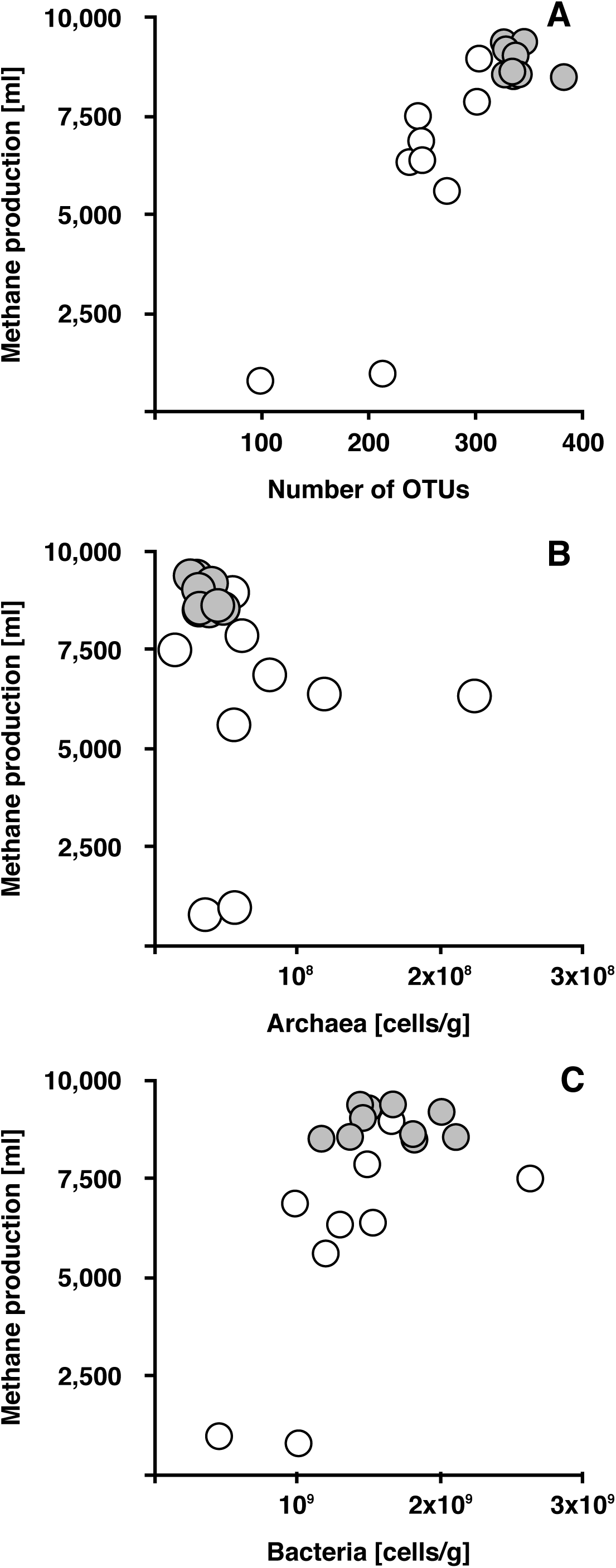
Within-community predictors of methane production. Relationships between A) Archaeal densities [cells/g] (*F*_1,15_ = 0.32, *P* > 0.2) B) Bacterial densities [cells/g] (*F*_1,16_ = 16.5, *P* < 0.001) and C) number of OTUs (*F*_1,16_ = 51.6, *P* < 0.001) and methane production [ml]. Note that qualitatively the same results apply when mixed communities are excluded from analyses.

The finding that coalescence results in the most productive individual community dominating the mixed community has direct implications for biotechnological uses of microbial communities. Given that the best performing community in isolation largely determined both the composition and performance of mixtures of communities, we hypothesized that methane production would increase with increasing number of communities in a mixture. We therefore inoculated laboratory-scale anaerobic digesters with 1, 2, 3, 4, 6 or 12 communities, ensuring that each of the 12 starting communities was only used once at each diversity level (see Extended Data Table 1). We found that cumulative methane production over a five-week period increased with increasing number of communities used as an inoculum (Figure 4). The positive correlation between community function and the number of inoculating communities is analogous to the commonly observed finding that community productivity increases with increasing species diversity (*20*). In our case, the mechanism underlying this positive relationship between the number of communities and productivity is a sampling effect: inoculating more communities increases the chance that the best performing community will be present in the mix. However, given that domination of the mixture by one community was not complete, it is possible that mixing communities could increase performance beyond that of the maximum of single communities in some circumstances.

**Figure 4:**
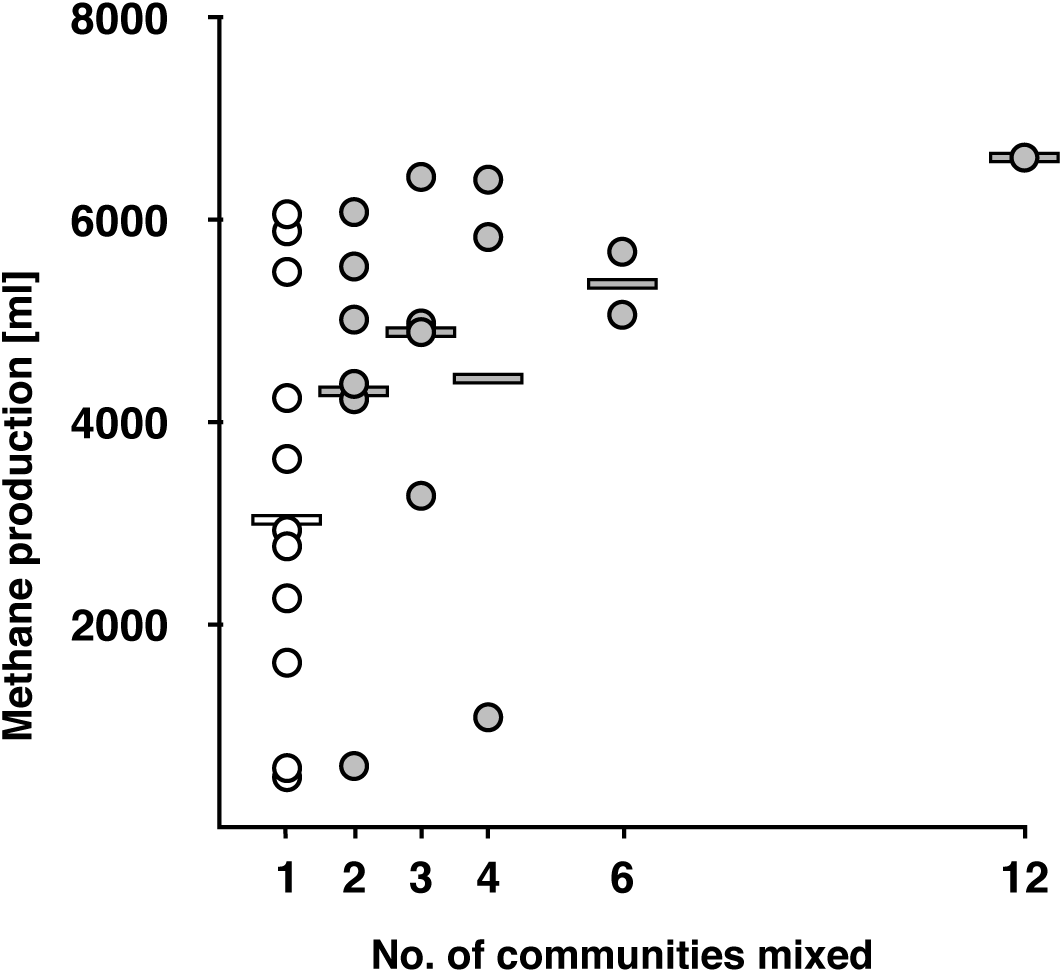
Cumulative methane production over time increases with number of inoculated communities. No single community was represented more than once at each diversity level, and there was a monotonic increase in methane production with number of communities used (*F*_1,26_ = 5.4, *P* = 0.03). Individual communities (white circles) and their average methane production (white line) are compared with mixes of communities (grey circles) and their averages (grey line) at different numbers of communities used.

Here, we have shown that coalescence of microbial communities results in dominance of a single community, and that the dominant community can be predicted from its original productivity.. These findings will have broad relevance to microbial communities, because the mechanism underlying the result relies on rules that underpin our understanding of community ecology: “niche-packed” communities use resources more fully and are harder to invade(*9*, *16*, *17*). Moreover, we have identified a way to significantly improve methane yield during anaerobic digestion: inoculate digesters with a broad range of microbial communities, and the best performing community will dominate. Given that resource use efficiency is often a desirable property of microbial communities, this approach could be applied to a range of biotechnological processes driven by microbial communities, as well as to manipulate microbiomes in clinical and agricultural contexts.

## Acknowledgements

The work was funded by the BBSRC, project BB/K003240/1. AB is additionally funded by the Royal Society, Axa Research Fund and NERC. KM & JH were funded through the project ENIGME from the INRA metaprogramme MEM (Meta-omics and microbial ecosystems). KM was additionally funded through an Institut Carnot 3BCAR international travel grant.

### Materials and Methods

#### Communities and fermentation

The communities used were collected from commercial operations, both from anaerobic digestors (AD plants) and communities present in nature used to seed the AD plants. The source and types of communities used can be found in Table 1. Communities were stored at 4°C prior to use. For all experiments, communities were grown in 500 ml bottles (600ml total volume with headspace; Duran) using the commercially available Automated Methane Potential Test System (AMPTS, Bioprocess Control Sweden AB) to measure CO_2_-stripped biogas production (referred to as methane in this paper). Samples were fed weekly in a fed-batch mode using a defined medium (see below for media composition). The communities from experiment 1 were equalised in terms of bacterial cells per gram of sample before inoculation using M9 salts to dilute them to the community with the lowest cell density, based on qPCR enumeration of 16S rRNA gene copies. For experiments 2 and 3, starting 16S rRNA copy number was determined (but not equalised between communities) and did not correlate with methane production. The fermenters were inoculated with 275 g of sample and fed with 25 ml of defined medium: meat extract 111.1 gl^-1^, cellulose 24.9 gl^-1^, starch 9.8 gl^-1^ glucose 0.89 gl^-1^, xylose 3.55 gl^-1^ (carbon to nitrogen ratio of 15:1) every week, starting with t_0_. Before the start of the fermentation, 0.3 mL of 1000x Trace Metal stock (1 gl^-1^ FeCl_2_. 4H_2_O, 0.5 gl^-1^ MnCl_2_. 4H_2_O, 0.3 gl^-1^ CoCl_2_. 4H_2_O, 0.2 gl^-1^ ZnCl_2_, 0.1 gl^-1^ NiSO_4_. 6H_2_O, 0.05 gl^-1^ Na_2_MoO_4_. 4H_2_O, 0.02 gl^-1^ H_3_BO_3_, 0.008 gl^-1^ Na_2_ WO_4_. 2H_2_O, 0.006 gl^-1^ Na_2_SeO_3_. 5H_2_O, 0.002 gl^-1^ CuCl_2_. 2H_2_O) was added to each fermenter. Experiment 1 (shown in Figure 1) ran for 5 weeks, experiment 2 (shown in Figure 2) for 6 weeks and experiment 3 (shown in Figure 4) for 5 weeks.

#### Measuring methane content of Biogas

All resulting lab-scale reactors inoculated with the samples were run at 37°C using the AMPTS. The AMPTS system measures the volume of biogas produced following stripping of CO_2_ from the produced gas. We have confirmed that the measured biogas is >95% methane using GC-FID.

#### DNA extraction, amplicon library construction and sequencing

DNA for 16S rRNA gene amplicon sequencing was extracted using QIAamp DNA Stool Mini Kit (QIAGEN) or FastDNA™ SPIN Kit for Soil (MP), depending on the experiment. Note that DNA extraction for mixed community P06 from the 10 mix experiment failed. The DNA for qPCR was extracted with the QIAamp DNA Stool Mini Kit (QIAGEN), protocol for pathogen detection with the 95ºC incubation step and the Powerlyzer Powersoil DNA KIT (MOBIO). DNA from *A. baylyi*, *P. fluorescens* SBW25 for Bacteria from *H. salinarum* DSM 669 for Archaea was used as standards. The primers(*21*) used to identify Bacteria were 16S rRNA 338f - ACT CCT ACG GGA GGC AGC AG, 518r - ATT ACC GCG GCT GCT GG for Archaea: 931f - AGG AAT TGG CGG GGG AGC A, m1100r - BGG GTC TCG CTC GTT RCC. The reagents used were: 1x Brilliant III Ultra-Fast SYBR^®^ Green QPCR Master Mix; 150nM 338f and 300nM 518r or 300nM 931f and 300 nM m1100r; ROX 300nM; and BSA 100 ng/µl final concentration. All samples were run in triplicate on a StepOnePlus (Applied Biosystems) qPCR machine using a program with 3’ 95ºC initial denaturation followed by 40 cycles of 5’’ at 95ºC and 10’’ at 60ºC, followed by a melting curve 95ºC for 15’’; 60ºC for 1’ ramping up to 95ºC in steps of 0.3ºC for 15’’ each. The melting curve analysis and the confirmation of the negative controls was done using Stepone Software v.2.3 (life technologies). The Cq values and the efficiencies of the samples and standards was determined as previously using LinRegPCR version 2016.0(*22*). The quantities were calculated using the one point calibration method as described earlier(*23*).

#### Analyses of sequenced samples

MiSeq amplicon reads were merged using Illumina-utils software(*24*). We quality-filtered only the mismatches in the overlapping region between read pairs using a minimum overlap (-- min-overlap-size) of 200 nt and a minimum quality Phred score (--min-qual-score) of Q20. We allowed no more than five mismatches per 100 nt (-P 0.05) over the 200 nt overlapping region.

Reads that fulfilled the quality criteria were analysed using Quantitative Insights Into Microbial Ecology (QIIME v.1.7)(*25*). We removed chimera using the *identify_chimeric_seqs.py* script, UCHIME reference ‘Gold’ database and USEARCH(*26*, *27*), which we also used to select OTUs. We assigned the taxonomy of our reads with QIIME *pick_open_reference_otus.py* function, using the Greengenes database version v13_8(*28*) as a reference with a minimum cluster size of 2 (i.e., each OTU must contain at least two sequences). We collapsed the technical replicates and filtered out the low abundance OTUs (<0.01% total, *filter_otus_from_otu_table.py*) and samples rarefied to an even depth of 26702 for both experiments where sequencing data is available. QIIME was used to calculate alpha and beta diversity data and produce NMDS plots.

For the NNLS analysis, following removal of low abundance OTUs and cumulative sum scaling transformation, the resulting .biom file was used to create a matrix 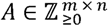(*m* rows of OTUs by *n* sample columns) for the seed bioreactors, and a column vector 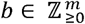 for each mixed bioreactor; both *A* and *b* hold non-negative integers of OTU abundances. One of the individual samples contained a negligible number of reads and was discarded from the analysis. The contribution, or weight, of each seed sample to the pattern of OTUs observed in a mixed sample is given by the column vector 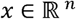 when solving for a system of linear equations *Ax* = *b*.

When modelling count data for environmental samples the fitted parameters of *x* will also be non-negative and the number of OTUs will usually exceed the number of samples (*m* > *n*). The task is to solve an over-determined system of linear equations where there are more equations than unknowns. It is likely that some of the linear equations will ‘disagree’ and there will be no exact solution. Geometrically, this may be interpreted as *b* not lying in the column space of *A*, a (hyper)plane holding the column vectors of *A*, or *Ax* − *b* ≠ 0. A least-squares approach may find the non-negative vector 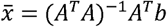 which is the projection of *b* back onto the column space of *A* that minimises the least-squares error ‘distance’ 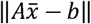. For our study the non-negative least-squares (NNLS) solution, 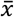, and least-squares errors were computed via the R packages ‘nnls’(*29*) and ‘limSolve’(*30*) for each of the mixed samples.

## References

1. R. MacArthur, E. Wilson, An equilibrium theory of insular zoogeography. Evolution (N. Y). 17, 373–387 (1963).

2. M. Vellend, Conceptual synthesis in community ecology. Q. Rev. Biol. 85, 183–206 (2010).

3. T. Kawecki, Adaptation to marginal habitats. Annu. Rev. Ecol. Evol. Syst. 39, 321–342 (2008).

4. M. Rillig, J. Antonovics, T. Caruso, A. Lehmann, Interchange of entire communities: microbial community coalescence. Trends Ecol. 30, 470–476 (2015).

5. M. Rillig, A. Lehmann, C. Aguilar-Trigueros, Soil microbes and community coalescence. Pedobiologia (Jena). 59, 37–40 (2016).

6. M. Gilpin, Community-level competition: asymmetrical dominance. Proc. Natl. Acad. Sci. U. S. A. 91, 3252–3254 (1994).

7. Y. Toquenaga, Historicity of a simple competition model. J. Theor. Biol. 187, 175–181 (1997).

8. C. Wright, Ecological community integration increases with added trophic complexity. Ecol. Complex. 5, 140–145 (2008).

9. M. Tikhonov, Community-level cohesion without cooperation. Elife. 5, e15747 (2016).

10. N. Hausmann, C. Hawkes, Plant neighborhood control of arbuscular mycorrhizal community composition. New Phytol. 183, 1188–1200 (2009).

11. G. Livingston, Y. Jiang, J. Fox, M. Leibold, The dynamics of community assembly under sudden mixing in experimental microcosms. Ecology. 94, 2898–2906 (2013).

12. C. Souffreau, B. Pecceu, C. Denis, An experimental analysis of species sorting and mass effects in freshwater bacterioplankton. Freshwater. 59, 2081–2095 (2014).

13. H. Adams, B. Crump, G. Kling, Metacommunity dynamics of bacteria in an arctic lake: the impact of species sorting and mass effects on bacterial production and biogeography. Front. Microbiol. 5 (2014).

14. R. MacArthur, Species packing and competitive equilibrium for many species. Theor. Popul. Biol. 1, 1–11 (1970).

15. J. Roughgarden, Resource partitioning among competing species—a coevolutionary approach. Theor. Popul. Biol. 9, 388–424 (1976).

16. A. Gardner, A. Grafen, Capturing the superorganism: a formal theory of group adaptation. J. Evol. Biol. 22, 659–671 (2009).

17. J. M. Levine, Species diversity and biological invasions: relating local process to community pattern. Science. 288, 852–4 (2000).

18. B. Schink, Energetics of syntrophic cooperation in methanogenic degradation. Microbiol. Mol. Biol. Rev. 61, 262–280 (1997).

19. K. Hillesland, D. Stahl, Rapid evolution of stability and productivity at the origin of a microbial mutualism. Proc. Natl. Acad. Sci. U. S. A. 107, 2124–2129 (2010).

20. D. Tilman, C. Lehman, Plant diversity and ecosystem productivity: theoretical considerations. Proc. Natl. Acad. Sci. U. S. A. 94, 1857–1861 (1997).

21. L. Øvreås, V. Torsvik, Microbial Diversity and Community Structure in Two Different Agricultural Soil Communities. Microb. Ecol. 36, 303–315 (1998).

22. J. M. Ruijter et al., Amplification efficiency: linking baseline and bias in the analysis of quantitative PCR data. Nucleic Acids Res. 37, e45 (2009).

23. R. Brankatschk, T. Fischer, M. Veste, J. Zeyer, Succession of N cycling processes in biological soil crusts on a Central European inland dune. FEMS Microbiol. Ecol. 83, 149–160 (2013).

24. A. M. Eren et al., Oligotyping: differentiating between closely related microbial taxa using 16S rRNA gene data. Methods Ecol. Evol. 4, 1111–1119 (2013).

25. J. G. Caporaso et al., QIIME allows analysis of high-throughput community sequencing data. Nat. Methods. 7, 335–336 (2010).

26. R. R. C. Edgar, Search and clustering orders of magnitude faster than BLAST. Bioinformatics. 26, 2460–2461 (2010).

27. R. C. Edgar, B. J. Haas, J. C. Clemente, C. Quince, R. Knight, UCHIME improves sensitivity and speed of chimera detection. Bioinformatics. 27, 2194–2200 (2011).

28. D. McDonald et al., An improved Greengenes taxonomy with explicit ranks for ecological and evolutionary analyses of bacteria and archaea. ISME J. 6, 610–8 (2012).

29. K. Mullen, I. Van Stokkum, The Lawson-Hanson algorithm for non-negative least squares (NNLS) (2007).

30. K. Soetaert, K. V. D. Meersche, D. V Oevelen, Package limSolve, solving linear inverse models in R. (2009).

